# Growing older, growing more diverse: sea turtles and epibiotic cyanobacteria

**DOI:** 10.1101/2024.03.08.584065

**Authors:** Lucija Kanjer, Klara Filek, Maja Mucko, Mateja Zekan Lupić, Maša Frleta-Valić, Romana Gračan, Sunčica Bosak

## Abstract

Cyanobacteria are known for forming associations with various animals, including sea turtles, yet our understanding of sea turtles associated cyanobacteria remains limited. This study aims to address this knowledge gap by investigating the diversity of cyanobacteria in biofilm samples from loggerhead sea turtle carapaces, utilizing a 16S rDNA amplicon sequencing approach. The predominant cyanobacterial order identified was *Nodosilineales*, with the genus *Rhodoploca* having highest relative abundance. Our results suggest that cyanobacterial communities became more diverse as sea turtles age as we had found a positive correlation between community diversity and the length of a sea turtle’s carapace. Since larger and older turtles predominantly utilize neritic habitats, the shift to more diverse cyanobacterial community aligned with a shift in loggerheads habitat. Our research provided detailed insights into the cyanobacterial communities associated with loggerhead sea turtles, establishing a foundation for future studies delving into this fascinating ecological relationship and its potential implications for sea turtle conservation.

## Introduction

Cyanobacteria form various symbiotic associations with a wide range of eukaryotic hosts including plants, fungi, sponges and microeukaryotes, each reaping distinct benefits. For instance, plant organisms and diatoms primarily form relationships utilizing cyanobacterial partner for nitrogen fixation, whereas animals and other heterotrophs benefit from the organic matter produced through photosynthesis (Adams et al. 2012). Organisms with simpler structures, such as sponges and corals, exhibit a tissue-embedded association with cyanobacteria, while associations with vertebrates are mostly superficial, occurring on the animal’s surface (Mutalipassi et al. 2021). It’s noteworthy that mutualistic connections between vertebrates and algae are relatively rare, with notable exceptions, such as the co-evolution of sloths with algae and cyanobacteria, resulting in a distinctive epizoic sloth associated photosynthetic community on their fur (Kaup et al. 2021).

Beside mutualistic and commensalistic association, cyanobacteria in aquatic environments have been observed to adversely impact animals, primarily due to their toxin production. However, the roles of benthic cyanobacteria remain understudied, especially in marine environment, as noted by Wood et al. (2020). Certain marine cyanobacteria have deterring effect on grazers and coral reef fishes due to their production of toxic compounds (Capper et al. 2016). In Egyptian sea waters, blooms of benthic cyanobacteria, specifically the *Oscillatoria acutissima* species, have been associated with significant fish mortality (Ismael 2012). Toxic cyanobacterial compounds have also been detected in dolphins (Brown et al. 2018, Davis et al. 2019). Cryptic cyanobacterial species were described on dolphins’ epidermis, though their impact on animal’s health remains unknown (A. O. Brown et al. 2021, J. Brown et al. 2021). Sea turtle health is also linked to cyanobacterial toxins. For instance, *Lyngbya majuscula*, a cyanobacterium growing epiphytically on sea grass consumed by green turtles, produces lyngbyatoxin, that has been associated with tumours and fibropapillomatosis in sea turtles(Arthur et al. 2008).

Sea turtles are large marine reptiles with seven species inhabiting the Earth today (Wyneken et al. 2013). Most of these species are endangered due to extensive array of human activities, including fishing bycatch, fishing gear ingestion, plastic pollution, and light pollution near nesting sites (Casale et al. 2018). Describing the animal-associated microbiota has been recognized as one of the supplementary measures for sea turtle conservation (Trevelline et al. 2019). Conservation biologists suggest that animal microbiomes research and the turtle health maintenance must become an integral part of conservation efforts, as it plays a crucial role in resolving conservation challenges such as native species reintroduction, captive breeding, non-native species invasion etc. (Redford et al. 2012, Peixoto et al. 2022). In recent years, there has been a growing effort to characterize and describe the endobiotic microbiota of the gut, cloaca and oral cavity of sea turtles (Abdelrhman et al. 2016, Biagi et al. 2019, Scheelings et al. 2020, Filek et al. 2021). However, the epibiotic microbiota of sea turtles, while frequently neglected, constitutes an equally vital component of the microbiome as the surface microbiome serves as the initial interface between an animal and its environment. The epibiotic communities are characterized by distinct factors like location, diet, and health status, which all contribute to the changes in the microbiome profile of these animals (Blasi et al. 2022, Kanjer et al. 2022).

While previous studies have documented cyanobacteria as part of the microbial biofilm growing on sea turtle carapaces(Blasi et al. 2022, Kanjer et al. 2022), a comprehensive exploration of the cyanobacterial community’s diversity is lacking. There is a notable absence of basic studies describing the “natural” cyanobacterial community in healthy sea turtles, making it challenging to discern differences between the cyanobacteria inhabiting sea turtles and those in other benthic habitats. The study of the cyanobacterial ecological diversity has traditionally employed various methods, ranging from microscopic determination to more recent molecular techniques. With the advancement of molecular biology, many new, up to now neglected cyanobacterial species and higher systematic ranks (orders, families) have been described, which could not be distinguished by morphology alone (Komárek 2020). Subsequently, cyanobacteriologists have adopted alternative methods to characterize biofilm cyanobacterial communities (Kolda 2018). Metabarcoding, i.e. amplicon sequencing, is one of the most commonly used techniques today, primarily due to its relatively low cost, proficiency, speed, established pipelines, and the absence of a requirement for specialized taxonomists or cultivation efforts (Kolda 2018). Recent efforts to enhance the accuracy of reference databases used for correct taxonomic identification (Roush et al. 2021, Lefler et al. 2023) offer a promising pathway for improving subsequent ecological studies.

This study aims to characterize the cyanobacterial community within the epibiotic biofilm on loggerhead sea turtle (*Caretta caretta*) carapaces, employing a metabarcoding approach through 16S rDNA amplicon sequencing. Additionally, we aim to explore variations in the cyanobacterial community composition across different sea turtle age groups, spanning from juveniles to sub-adults and adults. By doing so, this research aims to provide the initial comprehensive understanding of the diversity and composition of cyanobacteria linked to sea turtles, laying the groundwork for future investigations into host-associated cyanobacteria within marine environments.

## Methods

### Sampling procedures

Total of 28 loggerhead sea turtle carapaces were sampled in this study, of which 19 were juveniles, 5 sub-adults and 4 adults (**Table 1**). Turtles were classified as different developmental stage groups according to length of their carapaces and characterized as juveniles with curved carapace length (CCL) ≤59.9 cm, as subadults with CCL 60-69.9 cm and adults with CCL ≥70 cm according to Mariani et al. (2023) (**Figure 1**).

**Table 1.** Information about sampled loggerhead sea turtles; CCL = curved carapace length; CCW = curved carapace width. Turtle age groups were determined as follows: juveniles ≤59.9 cm CCL, sub-adults 60-69.9 cm CCL, adults ≥70 cm CCL.

| Sample ID | Turtle name/label | Age | CCL (cm) | CCW (cm) | Sampling Date DD.MM.YYYY | Admitted days before sampling | Rescue Centre |
| --- | --- | --- | --- | --- | --- | --- | --- |
| TB159 | Ella | Sub-adult | 62.0 | 58.0 | 30.06.2020 | 0 | Aquarium Pula |
| TB163 | Huanita | Sub-adult | 68.0 | 66.0 | 20.03.2020 | 7 | Aquarium Pula |
| TB167 | Maro | Sub-adult | 67.0 | 63.2 | 21.07.2020 | 26 | Aquarium Pula |
| TB175 | Maksimus | Juvenile | 45.2 | 44.2 | 28.07.2020 | 2 | Aquarium Pula |
| TB177 | Špela | Sub-adult | 68.0 | 67.0 | 3.8.2020 | 4 | Aquarium Pula |
| TB181 | CC_VIS_2001 | Juvenile | 42.0 | 37.0 | 29.06.2020 | 0 | Blue World Institute |
| TB183 | CC_VIS_2002 | Juvenile | 31.0 | 29.0 | 11.08.2020 | 0 | Blue World Institute |
| TB185 | CC_VIS_2003 | Juvenile | 31.0 | 28.0 | 20.08.2020 | 0 | Blue World Institute |
| TB189 | Valbiska | Juvenile | 30.0 | 27.5 | 5.11.2020 | 27 | Blue World Institute |
| TB191 | FS94 | Sub-adult | 67.5 | 66.0 | 30.11.2020 | 2 | Blue World Institute |
| TB195 | FS35 | Juvenile | 45.0 | 41.0 | 10.12.2020 | 12 | Blue World Institute |
| TB197 | FS25 | Adult | 70.0 | 67.0 | 19.12.2020 | 1 | Blue World Institute |
| TB199 | FS60 | Adult | 73.0 | 68.0 | 19.12.2020 | 1 | Blue World Institute |
| TB201 | Apox | Juvenile | 36.0 | 34.0 | 14.1.2021 | 2 | Blue World Institute |
| TB203 | CC_LOŠINJ_2102 | Juvenile | 55.0 | 53.5 | 25.01.2021 | 2 | Blue World Institute |
| TB205 | Zlata | Juvenile | 35.5 | 33.0 | 03.02.2021 | 4 | Blue World Institute |
| TB207 | Noemi | Juvenile | 40.5 | 38.0 | 16.02.2021 | 2 | Blue World Institute |
| TB209 | Sanjin | Juvenile | 32.0 | 29.0 | 16.02.2021 | 1 | Blue World Institute |
| TB211 | CC_LOŠINJ_2106 | Juvenile | 40.0 | 38.0 | 08.03.2021 | 1 | Blue World Institute |
| TB213 | Natanael | Juvenile | 33.0 | 29.0 | 10.03.2021 | 1 | Blue World Institute |
| TB215 | Karlo Albano | Adult | 73.0 | 73.0 | 25.03.2021 | 6 | Aquarium Pula |
| TB217 | Oliver Raul | Juvenile | 26.0 | 24.2 | 23.04.2021 | 3 | Aquarium Pula |
| TB219 | Martin | Juvenile | 37.8 | 34.8 | 23.04.2021 | 2 | Aquarium Pula |
| TB221 | CC_LOŠINJ_2112 | Juvenile | 31.0 | 28.0 | 25.05.2021 | 2 | Blue World Institute |
| TB223 | Bova | Adult | 70.0 | 64.0 | 08.06.2021 | 3 | Blue World Institute |
| TB227 | Calimero | Juvenile | 52.0 | 52.0 | 06.04.2021 | 5 | Blue World Institute |
| TB229 | Marijana | Juvenile | 26.0 | 23.0 | 06.04.2021 | 1 | Blue World Institute |
| TB231 | CC_LOŠINJ_2111 | Juvenile | 27.0 | 25.5 | 21.04.2021. | 1 | Blue World Institute |

**Figure 1.**
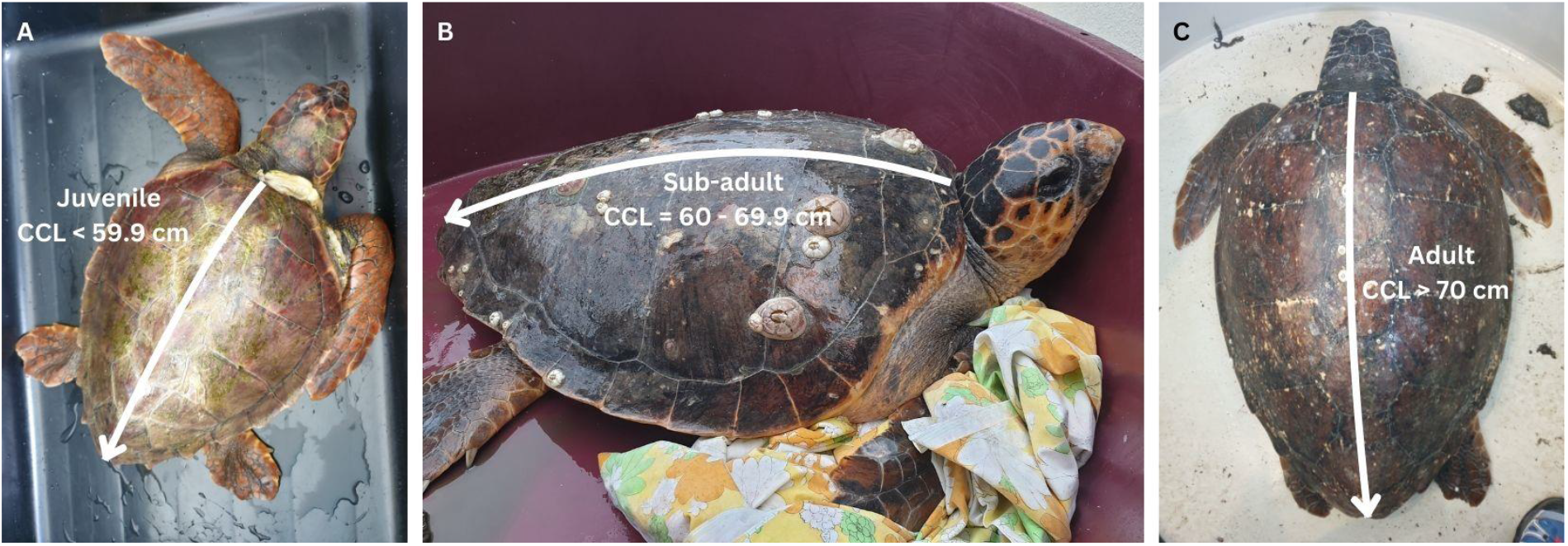
Examples of loggerhead sea turtles with indicated curved carapace length (CCL) measurements according to Mariani et al. (2023); A – juvenile sea turtle (TB189); sub-adult sea turtle (TB191); adult sea turtle (TB199).

Majority of animals that were sampled in this study were found stranded, injured, or were unintentionally caught by fishermen along the eastern coast of the Adriatic Sea in the period 2020-2021 (**Figure 2**) and transported to two Sea Turtle Rescue Centre’s in Croatia, to Blue World Institute in Mali Lošinj and to Aquarium Pula in Pula. Three turtles (TB181, TB183 and TB185) were sampled during research activities carried out by the Blue World Institute in Vis archipelago, Croatia, and were immediately released (**Table 1**). Sampling was conducted by trained personnel in accordance with the 1975 Declaration of Helsinki, as revised in 2013 and the applicable national laws, as soon as possible upon the turtles’ arrival to the rescue centres. The photographs of animals included in the study are available in the Supplement (Figure S2). A non-invasive approach was used for acquiring microbial biofilm from carapace. The epibiotic microbial biofilm was randomly collected from the entire area of the turtle carapaces by brushing the surface with a clean toothbrush (Dentalux) and resuspended in 96% ethanol in 50 ml collection tubes. In addition, we collected two samples of microbial biofilm growing on the plastic surfaces of the inner side of rehabilitation pools where turtles TB217 (sample ID: TB217P) and TB219 (sample ID: TB219P) were housed as well as a single sample of scrapings obtained from several submerged rocks (sample ID: VRS) collected at the beach near Aquarium Pula (44.833372, 13.833015). All collected samples were kept in 96% ethanol at -20 °C prior to DNA extraction.

**Figure 2.**
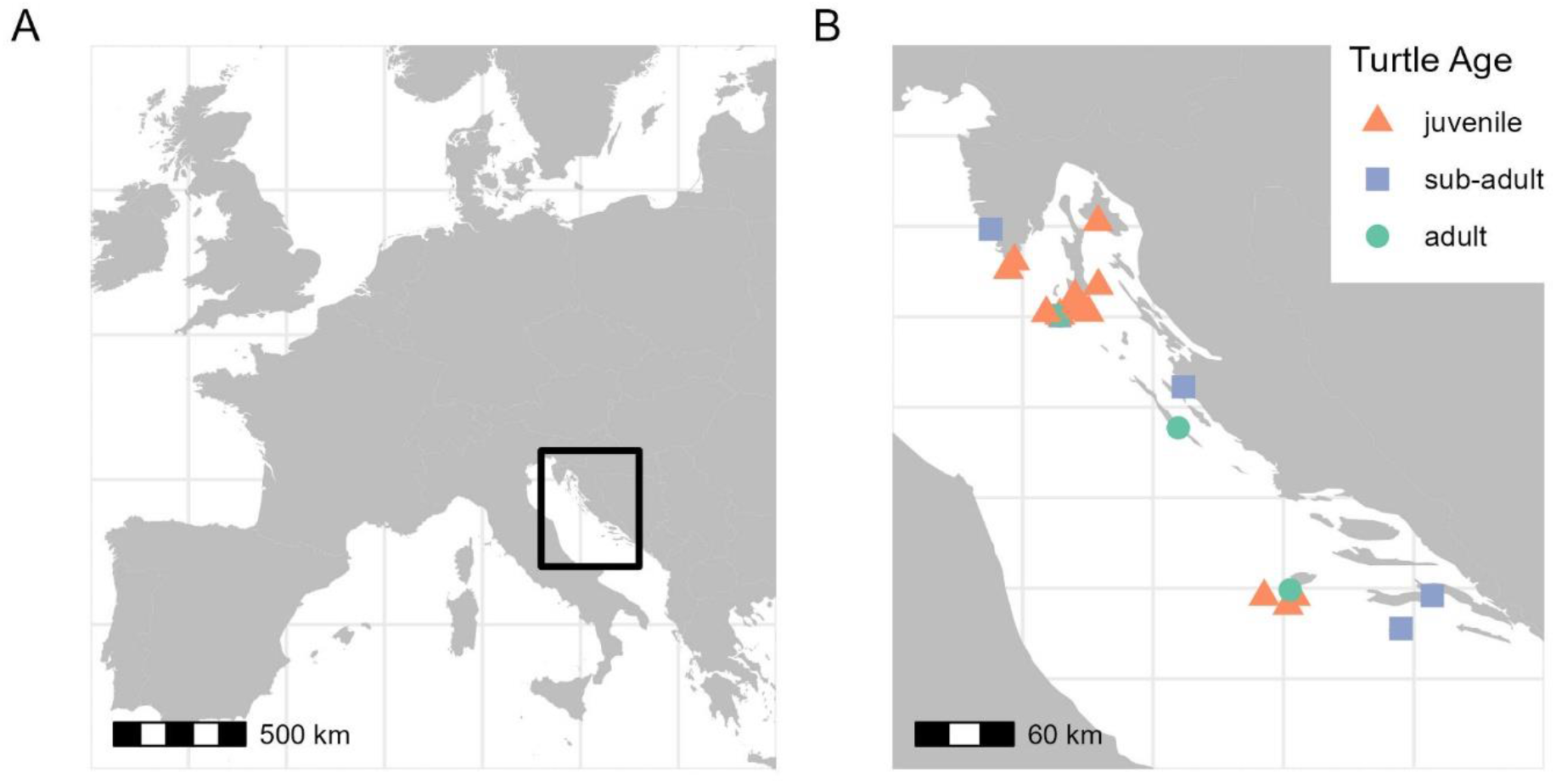
Maps of locations of sampled loggerhead sea turtles; A – map of Europe with indicated study area in black rectangle; B – map of Adriatic Sea with indicated locations of each sampled animal, triangle shapes represent juvenile animals, squares represent sub-adult animals and circles represent adult animals.

Scanning electron microscopy observations of several scutes shed from the carapaces from turtle TB235 (named Kolumbo, CCL = 65 cm, sampled by Sea Turtle Rescue Centre in Aquarium Pula on 13.10.2023) was performed to visualize the cyanobacteria as part of the epibiotic microbial biofilm. This sample was only used for morphological analyses and not included in amplicon sequencing analysis. The scutes were firstly chemically dried with HMDS (Oshel 1997). The small pieces of the scutes were then mounted on aluminium stubs with carbon tape and sputter-coated with platinum (15 nm) using a Precision Etching and Coating System, PECS II (Gatan Inc., Pleasanton, CA, USA). The samples were analysed with scanning electron microscope JEOL JSM-7800F (JEOL, Tokyo, Japan) at the Department of Physics, Centre for Micro and Nano Sciences and Technologies, University of Rijeka, Croatia.

### DNA processing

Prior to DNA extraction, the samples were centrifuged in 15 ml tubes (2500 rpm for 10 min, Centurion Scientific K3 Series, UK) and ethanol in supernatant was removed leaving only pelleted sample which was used for downstream analyses. DNA was isolated from 250 mg of sample using DNeasy PowerLyzer PowerSoil extraction kit (Qiagen, Germany). Manufacturers guidelines were followed with a few modifications described below. After adding the C1 solution, samples were incubated at 70 °C for 10 min. For bead beating Qiagen TissueLyser (Retsch, Germany) was used at 30 Hz for 1 min. In the end DNA was eluted in 50 μl of Solution C6 with 5 min incubation at room temperature before final centrifugation step. Nuclease-free water (W4502 Sigma-Aldrich) was used as the negative control and was processed using the DNA extraction kit in parallel to all the samples. The quantity and purity of DNA was measured on Biospec-nano spectrophotometer (Shimadzu, Japan). Samples of extracted DNA were sent for library preparation and sequencing in Microsynth AG (Balgach, Switzerland). Sequencing was performed on Illumina MiSeq Systems platform generating 2×300 bp pair-end reads using cyanobacteria-specific custom primers targeting V6 region: 1328F (5’GCTAACGCGTTAAGTATCCCGCCTGG-3’) and 1664R (5’-GTCTCTCTAGAGTGCCCAACTTAATG-3’) (Lee et al. 2017).

### Sequence data analysis

Sequences were trimmed from primers and adapters and demultiplexed by sequencing agency. Demultiplexed sequences were imported into QIIME2 environment, version 2022.8 (Bolyen et al. 2019) using Casava 1.8 demultiplexed paired-end format. Sequences were denoised using DADA2 (q2-dada2 plugin) to amplicon sequence variants (ASV) (Callahan et al. 2016). V6 sequences were truncated at 258 bp for forward reads and 162 bp for reverse reads. V6 sequences were truncated at 240 bp for forward reads and 200 bp for reverse reads. Taxonomy was assigned to ASVs using q2-feature-classifier plugin (Bokulich et al. 2018). CyanoSeq reference database incorporated into SILVA v.138 was used to assign taxonomy (Lefler et al. 2023). Prior to downstream analyses, chloroplast sequences were excluded from dataset, leaving only phylum *Cyanobacteriota* for subsequent analysis. QIIME2 artifacts table, taxonomy, rooted tree and metadata table were imported to R environment (R version 4.2.2, R Studio version 2023.09.1) using qime2R package (Bisanz 2018) and further analysed using phyloseq (McMurdie and Holmes 2013) and vegan (Oksanen et al. 2020) packages while visualizations were made by ggplot2 (Wickham 2016) package. Cyanobacterial community composition on Order and Genus level was illustrated using compositional taxa barplots. For alpha and beta diversity analyses, we rarefied dataset to the read depth of 10,098. Due to low number of reads (less than 10,098) samples TB211, TB213, TB167, TB217P, TB219P and negative control were excluded from diversity analyses. Two alpha diversity metrices were calculated: observed ASV richness and Shannon diversity index. The Pearson correlation test was used to calculate the relationship between alpha diversity metrics and curved carapace length (CCL). Beta diversity was estimated using robust Aitchison distance and visualized using principal components analysis (PCA) for robust Aitchison distance on center log ratio (CLR) transformed data using q2-deicode plugin (Martino et al. 2019). The permutational multivariate analysis of variance (PERMANOVA) was used to test for significant differences between sample groups based on estimated turtle age (“juvenile”, “sub-adult” and “adult”). Map was made in R using additional packages sf (Pebesma 2018, Pebesma and Bivand 2023), rnaturalearth (South et al. 2023), rnaturalearthdata (South et al. 2024) ang ggspatial (Dunnington et al. 2023). Images are edited using Canva editor.

## Results

### Community composition

Scanning electronic microscopy analyses of the biofilm formed on loggerhead scutes showed presence of homocytous filamentous cyanobacterial trichomes (**Figure 3**) embedded with organic matter and other prokaryotic and microeukaryotic cells. They most likely belong to orders like *Oscillatoriales* (**Figure 3A, B**), *Spirulinales* (**Figure 3C, D**), *Nodosilineales, Leptolyngbyales, Oculatellales* (**Figure 3E**), *Pseudanabenales* or *Prochlorotrichales* (**Figure 3F**), however based solely on morphological characteristics, exact taxonomical identification was not possible.

**Figure 3.**
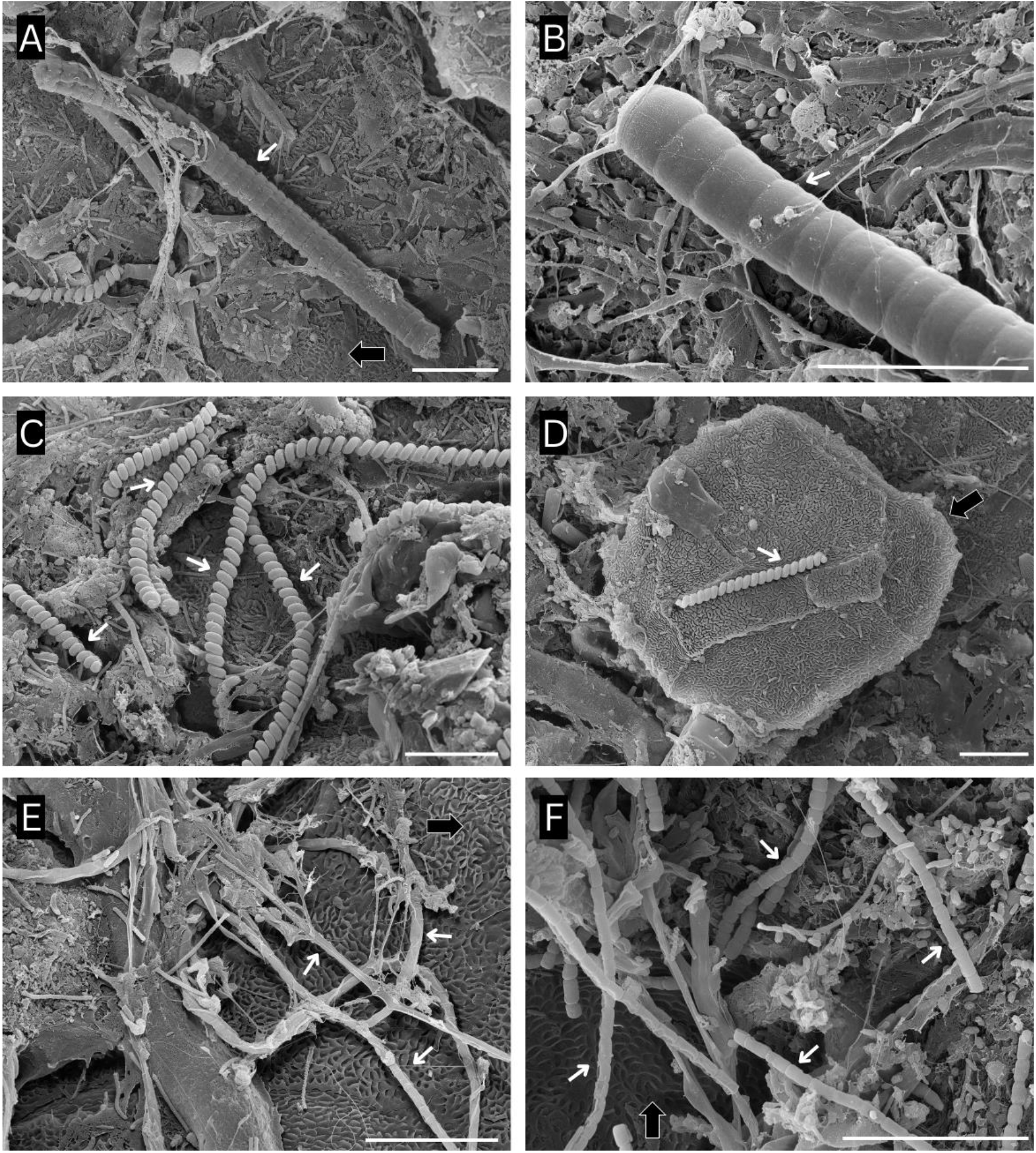
Scanning electron microscopy (SEM) images of cyanobacteria (white arrows) in the microbial community growing on the sub-adult loggerhead sea turtle carapace surface (black arrows) sampled from sea turtle TB235; *Oscillatoriales* (A, B); *Spirulinales* (C, D); *Nodosilineales, Leptolyngbyales* or *Oculatellales* (E); *Pseudanabenales* or *Prochlorotrichales* (F); Scale bar represents length of 10 μm.

High throughput sequencing yielded 2,359,741 high quality cyanobacterial sequences arranged in 551 ASVs and in 30 samples. Median sequence frequency per sample is 20,069.5 (maximum 412,998.0, minimum 107.0). In sample TB211 no cyanobacterial sequences were identified and therefore the sample was excluded from further analyses.

The predominant cyanobacterial order in whole dataset was *Nodosilineales*, followed by *Prochlorotrichales* and *Chrococcales* (**Figure 4**). On genus level *Rhodoploca, Leptothoe* and *Cymatolege* had highest relative abundance.

**Figure 4.**
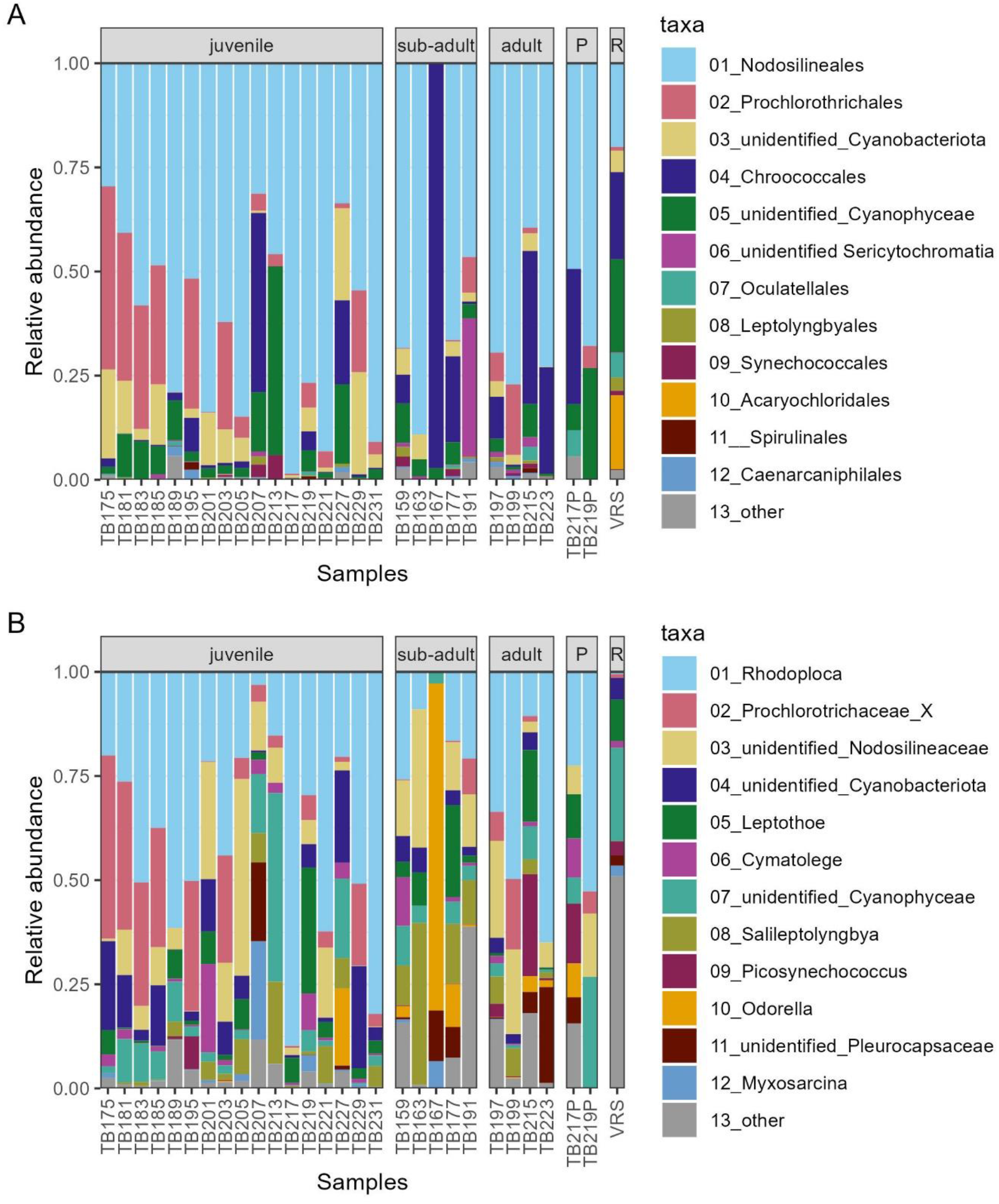
Relative abundances of 12 orders (A) and genera (B) with highest relative abundance identified with CyanoSeq database, remaining orders (A) and genera (B) were put in category “other” for better visualisation. One bar represents individual turtle carapace samples organized according to age class; P = pool sample and R = rocks sample.

There were nine core cyanobacteria features (ASVs) identified in 100% of samples in adult sea turtle group (**Table S1**) and they were identified as belonging to genera *Rhodoploca, Salileptolyngbya, Picosynechococcus* and *Odorella*. There were also four core ASVs that were identified only to the family level *Nodosilineaceae* (BLAST showed 98.45% identity with *Leptolyngbya* sp. PCC 7124), phylum *Cyanobacteriota* (BLAST showed 100% identity with *Cyanobacterium* sp. BBD-AO-Red), order *Caenarcaniphilales* (BLAST showed 99.61% identity with Uncultured bacterium clone QSW25) and class *Cyanophyceae* (BLAST showed 98.06% identity with *Copelandiella yellowstonensis* YNP83A-MA6 and *Geitleribactron purpureum* Tovel-1). Two core ASVs were present in 100% samples in the juvenile turtle group (**Table S2**), one belonged to the genus *Rhodoploca*, and the other ASV as belonging to the family *Nodosilineaceae* (BLAST showed 98.45% identity with *Leptolyngbya* sp. PCC 7124). In sub-adult class there were no core features identified in 100% of samples, but there were five features identified in 80% of samples (**Table S3**). Those cyanobacteria belong to the genera *Rhodoploca, Salileptolyngbya, Odorella*, one ASV belonging to the family *Nodosilineaceae* (BLAST showed 98.45% identity with *Leptolyngbya* sp. PCC 7124), one belonging to class *Cyanophyceae* (BLAST showed 98.06% identity with *Copelandiella yellowstonensis* YNP83A-MA6 and *Geitleribactron purpureum* Tovel-1) and one ASV belonging to phylum *Cyanobacteriota* (BLAST showed 100% identity with *Cyanobacterium* sp. BBD-AO-Red).

In pool sample TB217P the predominant cyanobacterial order was also *Nodosilineales*, but in contrast to carapace of juvenile TB217, pool sample had also high relative abundance of orders *Chroococcales* and *Oculatellales* (**Figure 4A**). Regarding identified genera, on turtle TB217, the most dominant was *Rhodoploca*, while in the pool sample there was mix of *Rhodoploca, Leptothoe, Cymatolege, Picosynechococcus* and *Oderella* (**Figure 4B**). Furthermore, pool sample TB219P and carapace biofilm from juvenile TB219 were predominated by order *Nodosilineales* (**Figure 4A**). On genus level, TB219 had high relative abundance of *Rhodoploca, Leptothoe* and *Cymatolege*, while pool sample TB219P was dominated by *Rhodoploca* and other unidentified cyanobacteria (**Figure 4B**).

Rock scraping sample VRS was not dominated by single order, rather a mixture of highly abundant *Nodosilineales, Chroococcales, Oculatellales* and *Acaryochloridales*, of which last was not recorded as prevalent in any of the turtle samples (**Figure 4A**).

### Alpha diversity

The richness and diversity of cyanobacterial communities in each sample was estimated using observed ASVs and Shannon diversity index, respectively (**Figure 5A** and **Figure 5B**). There were between 27 and 196 observed cyanobacterial ASVs with median value of 47 ASVs (**Table S4**). Turtles’ carapace size and observed ASV richness were positively correlated (Pearson correlation R = 0.449), but not statistically significant (p-value > 0.05; **Figure 5A**). Shannon diversity index ranged between 0.619 and 3.824 with median value of 1.97 (**Table S4**). Turtles’ carapace size and Shannon diversity index were significantly positively correlated (Pearson correlation R = 0.699; p-value = 0.0009; **Figure 5B**).

**Figure 5.**
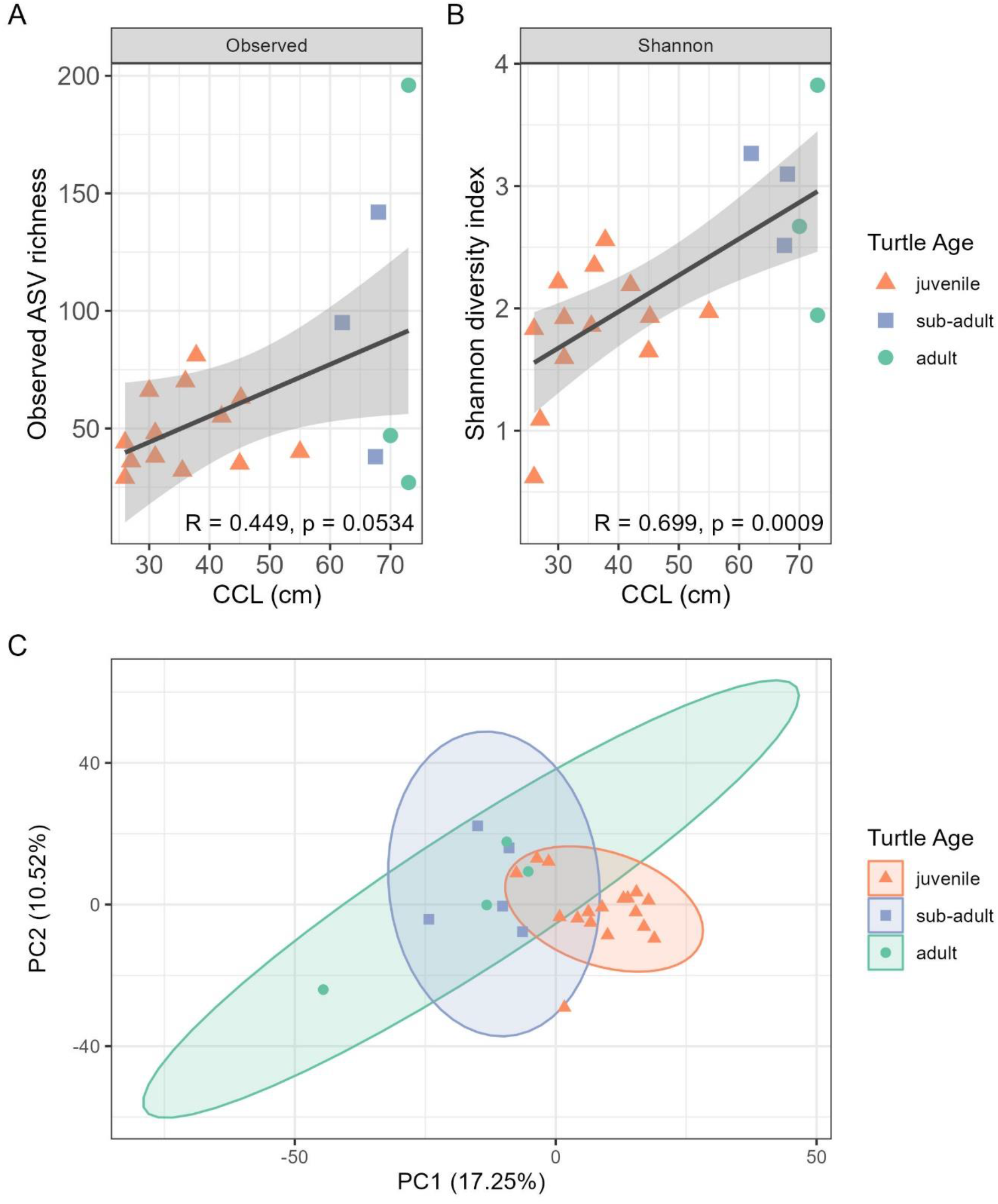
Alpha and beta diversity visualizations; observed amplicon sequence variant (ASV) richness compared to curved carapace length (CCL), juveniles n = 13, sub-adults n = 3, adults n = 3 (A); Shannon diversity index compared to CCL, juveniles n = 13, sub-adults n = 3, adults n = 3 (B); principal components analysis (PCA) based on central log (CLR) transformed data, juveniles n = 18, sub-adults n = 5, adults n = 4 (C); triangle shapes represent juvenile animals, squares represent sub-adult animals and circles represent adult animals.

### Beta diversity

Principal Components Analyses of robust Aitchison distance (rPCA) between three age classes (26 samples, **Figure 5C**) and between all groups (29 samples, **Figure S2**) indicate groupings based on estimated turtle age where samples of juvenile turtles cluster tightly together, while samples of adult and sub-adult turtles are grouped together and adult samples span over both sub-adults and juveniles (Figure 3C). PERMANOVA results (**Table 2**) showed significant differences in cyanobacterial community based on estimated turtle age class. Pairwise PERMANOVA showed significant difference between cyanobacterial communities sampled from both adult and sub-adult versus juvenile sea turtles (**Table 2**). However, there was no significant difference between cyanobacterial communities of adult and sub-adult sea turtles (**Table 2**).

**Table 2.** Permutational multivariate analysis of variance (PERMANOVA) on robust Aitchison distance of CLR transformed data of cyanobacterial abundances in samples of sea turtle biofilm based on turtle developmental stage (groups “juvenile” (n = 18), “sub-adult” (n = 5) and “adult” (n = 4)); done based on 999 permutations; asterisk (*) indicates significant difference between groups (p-value and q-value < 0.05).

| Group 1 | Group 2 | pseudo-F | p-value | q-value |
| --- | --- | --- | --- | --- |
| adult | juvenile | 3.398 | **0.036\*** | 0.108 |
| adult | sub-adult | 0.932 | 0.477 | 0.716 |
| juvenile | sub-adult | 6.496 | **0.001\*** | **0.006\*** |

## Discussion

This study offers a first overview of the cyanobacterial diversity within the microbial epibiotic communities of loggerhead sea turtles inhabiting the Mediterranean Sea. Our findings demonstrate the prevalence of the order *Nodosilineales* in all sea turtle samples, with the genus *Rhodoploca* emerging with highest relative abundance. Our results show a noticeable increase in cyanobacterial community diversity associated with transition from juvenile to adult sea turtles, since adult stages expand their migratory routes to breeding and nesting grounds. This suggests a marked shift in cyanobacterial composition throughout the life stages of loggerhead sea turtles, shedding light on the dynamic nature of their microbiota.

In our epibiotic biofilm samples, the predominant order was *Nodosilineales*, a recently established order in the revised cyanobacterial taxonomy (Strunecký et al. 2023). Cyanobacteria belonging to the order *Nodosilineales* are primarily filamentous, with cell width below 2 μm and parietal thylakoids, exhibiting a highly diverse ecological distribution, ranging from marine to freshwater environments (Strunecký et al. 2023). More specifically, ASVs belonging to the order *Nodosilineales*, the family *Cymatolegaceae*, represented by the genera *Rhodoploca, Cymatolege*, and *Leptothoe*, were highly prevalent in our turtle epibiotic samples. *Rhodoploca* was not only the most prevalent genera, but also the core taxa in juvenile, sub-adult and adult turtles of this study. *Cymatolegaceae* species are predominantly marine or brackish, displaying variability in cell morphology, with the pink trichomes as their shared distinctive feature. Notably, all three genera were isolated and described from marine sponges in the Aegean Sea (Caroppo et al. 2012, Konstantinou et al. 2019, 2021). This group also exhibits variability in the photosynthetic apparatus, with some purportedly lacking chromatic adaptation or utilizing aforementioned far-red light for photosynthesis (Strunecký et al. 2023).

Second most dominant order, *Prochlorotrichales*, represented so far with one genus *Prochlorothrix* (Strunecký et al. 2023) was well represented within juvenile turtles, but lacking in sub-adults and adults apart from TB191 and TB199, respectively (**Figure 4A**). *Prochlorotrichales* with non-motile trichomes in contrast to sheath-bearing motile *Nodosilineales* could be in disadvantage living on turtle carapace. This could explain their higher abundance in less diverse communities of juvenile individuals prior to *Nodosilineales* predominant niche occupation. One of the ASVs identified as core taxa (*Cyanobacterium* sp. BBD-AO-Red) is found to be connected to black band disease of corals worldwide (Buerger et al. 2016). Another core ASV was identified either as *Copelandiella yellostonensis* (Kaštovský et al. 2023), known as mat forming thermophilic cyanobacterium cryptic to *Leptolyngbya* or *Geitleribactron purpureum*, unicellular and heteropolar cyanobacterium that forms purple biofilms on rocks (Cantonati et al. 2014) from *Leptolyngbyaceae* family (Mareš and Cantonati 2016).

The utilization of far-red light is commonly observed in cyanobacteria residing in conditions with low light, representing a survival strategy in otherwise challenging environments (Gan et al. 2014, Antonaru et al. 2020). The dominant cyanobacterial orders in this study, *Nodosilineales* and *Prochlorotrichales*, are known as cyanobacteria capable of far-red light photoacclimation (FaRLiP). It provides the cells fast adaptation mechanism which can be general advantage when living on a moving and diving host (Gisriel et al. 2023). However, the presence of far-red light utilization is a trait that may or may not be exhibited by members of the same taxonomic group and cannot be conclusively confirmed based solely on the 16S rRNA gene (Antonaru et al. 2020). While this research does not delve into testing the real presence and relative abundance of far-red light-utilizing cyanobacteria, it is important to note that such utilization could enhance survival in habitats like the sea turtle carapace, where light exposure often fluctuate due to the host animal’s diving behaviour. This is especially pronounced during winter when turtles may express overwintering behaviour and enter into dormant state on the sea bottom with occasional quick excursions to the surface (Hochscheid et al. 2005).

The main characteristic of the eastern Adriatic coastal environment is its limestone rocky shore, with a very limited number of sandy beaches. As a result, previous research has primarily focused on endolithic and supratidal cyanobacteria (Brandes et al. 2015, Palinska et al. 2017, Vondrášková et al. 2017, Vogt et al. 2019). The most dominant cyanobacteria identified in these studies belonged to the former order *Pleurocapsales*, notorious order of morphologically heterogenous cyanobacteria having random baeocyte formation throughout younger pseudofilaments (Komárek et al. 2014), which is now classified as the family *Pleurocapsaceae* under the order *Chrococcales* (Strunecký et al. 2023).

These cyanobacteria are commonly found in epilithic and endolithic habitats worldwide. A study by Kolda et al. (2020) compared cyanobacteria in sediment and coastal water in habitats under anthropogenic pressure in Croatia and found that the family *Pleurocapsaceae* was dominant, primarily the genera *Xennococcus* and *Pleurocapsa*. In our turtle epibiotic samples, the order *Chrococcales* and family *Pleurocapsaceae* had notable relative abundance, but they were not dominant. The genus *Odorella*, which was described recently as a cryptic genus to *Pleurocapsa* (Shalygin et al. 2019), ranks high among identified genera and it is possible that previous study by Kolda et al. (2020) may have identified it as *Pleurocapsa*. Notably, *Odorella* is the first genus in the family *Pleurocapsaceae* found to produce microcystins and various other cyanotoxins including 38 bioactive molecules in total. *Odorella* and unidentified *Pleurocapsaceae* were more common in sub-adult and adult turtle samples. In terms of diversity and taxonomic richness, our samples exhibited higher Shannon diversity compared to cyanobacterial communities obtained from tidal flats in Croatia (sample ASV richness 53, Shannon diversity 0.8), but were found to be less rich in amplicon sequence variants (ASVs) (median = 47) (Vogt et al. 2019). Another study on supratidal endolithic cyanobacteria (Brandes et al. 2015) showed lower richness (OTU richness 2-25) but higher diversity (average Shannon diversity 4.13) than the samples from our study.

In understanding the ecological relationship between cyanobacteria and turtles, it is essential to draw parallels with already well-studied associations. For instance, studies on diatoms suggest a significant level of host specificity, where they are consistently found on all sea turtles, and the community is predominantly comprised of putatively epizoic species (Robinson et al. 2016, Ashworth et al. 2021). Unlike diatoms, cyanobacteria do not exhibit universal presence on all sampled turtles (Kanjer et al. 2022), with some animals not hosting any cyanobacterial sequences. Previously, Blasi et al. (2022) also reported small proportion of cyanobacteria on *C. caretta* carapace (less than 1% of whole community), identifying *Pseudoanabenaceae* and *Rivulariaceae* as inhabitants of mainly anterior scutes, suggesting a less competitive environment for Cyanobacteria in respect to more diverse and prominent macroalgae. Despite a substantial portion of unidentified sequences, our findings suggest that cyanobacteria inhabiting sea turtles are not inherently host-specific. Instead, the turtle’s carapace appears to carry variety of common benthic taxa, some found in sea sponges, like family *Cymatolegaceae*, that are known to be capable of adapting to diverse light conditions and also to produce various metabolically active substances, like cyanotoxins.

The life cycle of the loggerhead sea turtle can be delineated into four phases: hatchling, juvenile, sub-adult, and adult. Throughout their life cycle, they undergo an ontogenetic habitat shift from the oceanic to the neritic phase (Wyneken et al. 2013). Hatchling and juvenile turtles are in the oceanic phase, wherein they typically swim and feed in open waters due to the underdevelopment of their lungs, preventing them from diving deep for feeding. As their lungs mature, they gain the ability to dive deeper and feed in benthic habitats closer to the coast, a distinctive feature of their sub-adult phase and the transition from oceanic to neritic habitats. In their sexually active adult stage, they predominantly inhabit and feed in benthic environments during their neritic phase. Adult loggerheads are omnivores and known as bioturbators of marine sediments during their foraging behaviour and thus increase the chances of new cyanobacteria colonisation on their carapace (Lazar et al. 2011). Recent studies (Baldi et al. 2023, Mariani et al. 2023) have indicated that in the Mediterranean loggerhead population, the distinction between the oceanic and neritic phases is less pronounced, primarily because of easily accessible shallow foraging grounds, particularly in the Adriatic Sea (Casale et al. 2008). Nevertheless, our results reveal clear differences in cyanobacterial communities between adults and juveniles, reflecting certain aspects of turtle biology and ecology.

During the juvenile phase, carapace growth is accelerated (Casale et al. 2011) leading to more frequent shedding of old cyanobacterial biofilm and replacement with new scutes. Since formation of biofilm on various types of surfaces is initialized with bacteria and cyanobacteria and it is extremely fast process (within hours), this dynamic could create additional microenvironments for cyanobacteria to colonize (Rubio 2002, Mejdandžić et al. 2015, Dang and Lovell 2016, Grzegorczyk et al. 2018). The observation that juvenile turtles exhibit less rich and diverse cyanobacterial communities can be possibly explained by their less frequent contact with benthic communities and repetitive primary biofilm formation compared to adult turtles which have more diverse communities build from different stages of biofilm formation. Although juveniles in the Mediterranean may feed in the same locations as adults, it does not imply that they spend the same amount of time in these locations, potentially resulting in a disparity in cyanobacterial exposure. Furthermore, another plausible explanation lies in the ecology of microorganism itself, where the age of the biofilm is correlated with more diverse and layered communities (Fenchel and Kühl 2000, Mejdandžić et al. 2015).

Recent studies suggest that microcystin production might be more prevalent among marine cyanobacteria than previously believed (Konstantinou et al. 2019). Microcystin, a cyanotoxin produced by certain cyanobacteria, has demonstrated antibiotic properties in various contexts. This unique attribute suggests potential applications for microcystin in the development of novel antibiotics or antimicrobial agents, opening avenues for further exploration in the field of medical research. While microcystin production in sea turtle biofilm could be advantageous in a way where it acts as a natural deterrent against pathogenic bacteria growth (Ramos et al. 2015), it also raises concerns about potential toxicity to vertebrates (Zhou et al. 2021), including the host turtle. The close association of toxin-producing cyanobacteria with sea turtles could pose an even greater threat to these already endangered animals. The cyanobacterial diversity in juvenile turtles is particularly susceptible to the dominance of a single species. Given that cyanotoxins can be present in taxa residing on sea turtles, there is a possibility that the turtles’ carapace is infiltrated by toxic cyanobacteria, potentially contributing to their compromised health and subsequent hospitalization. Notably, the associations between cyanobacteria and sea turtles have not been previously studied, and these factors were not considered in the admission criteria for turtles in rescue centres.

## Conclusion

Our research provides the first detailed overview of the cyanobacterial community associated with loggerhead sea turtles, revealing a shift in cyanobacterial diversity as sea turtles age. This raises new questions and possibilities in cyanobacterial research and also enhances our understanding of sea turtle biology and ecology. Given the high number of sequences that could not be identified, we recommend increased culturing efforts directly from sea turtle biofilm to elucidate this complex picture. Furthermore, investigating sea turtle microbiome as potential sources of cyanobacterial strains with far-red acclimation presents a captivating avenue for ecological research. In conclusion, we propose the implementation of cyanotoxin screening in the diagnostic processes of hospitalized sea turtles, particularly when the cause of the turtle’s compromised health is not immediately evident.

## Data Accessibility

Demultiplexed sequences are available at EMBL ENA database under accession number PRJEB68310. Code, metadata, input files, instructions and visualizations for analyses of sequence data are available via GitHub on link https://github.com/lucijakanjer/turtle-cyano-NGS.

## Abbreviations

ASV: amplicon sequence variant
CCL: curved carapace length
CCW: curved carapace width
CLR: center log ratio
PERMANOVA: permutational multivariate analysis of variance

## Acknowledgements

We are thankful to Dr. Milena Mičić, Karin Spiech, Simona Matas, DMV and the rest of the staff from the Marine Turtle Rescue Center (Aquarium Pula) as well as Dr. Draško Holcer and Tina Belaj, DMV (Blue World Institute) for their help in sample collection and to Prof. Zlatko Liber for providing the laboratory space and equipment for DNA isolations. We also thank to dr. Ivana Jelovica Badovinac (Faculty of Physics, University of Rijeka) for the help with SEM imaging. Another thanks to Forrest W. Lefler for providing CyanoSeq sequences and taxonomy files for input in QIIME2. This study was supported by the Croatian Science Foundation under the project number UIP-2017-05-5635 (Loggerhead sea turtle (*Caretta caretta*) microbiome: Insight into endozoic and epizoic communities – TurtleBIOME).

## Authors contribution

Lucija Kanjer: conceptualization, formal analysis, investigation, methodology, visualization, writing – original draft preparation; Klara Filek: investigation, methodology, writing – review & editing; Maja Mucko: methodology, writing – review & editing; Mateja Zekan Lupić: resources, writing – review & editing; Maša Frleta-Valić: resources, writing – review & editing; Romana Gračan: conceptualization, project administration, writing – review & editing; Sunčica Bosak: conceptualization, funding acquisition, project administration, supervision, writing – review & editing.

## Supplementary material

**Table S1**. List of core ASV features found on 100% sampled adult sea turtles.

**Table S2**. List of core ASV features found on 80% sampled sub-adult sea turtles.

**Table S3**. List of core ASV features found on 100% sampled juvenile sea turtles.

**Table S4**. Alpha diversity indices.

**Figure S1**. Photographs of sampled loggerhead sea turtles taken by Blue World Institute and Sea Turtle Rescue Centre Aquarium Pula; photographs of animals TB163, TB175, TB177, TB217, TB219, TB227, TB229 and TB231 were not taken.

**Figure S2**. PCA of center log ratio (CLR) transformed abundances for all samples.

