## Supplements for "Growing older, growing more diverse: sea turtles and epibiotic cyanobacteria": Figure S1.pdf

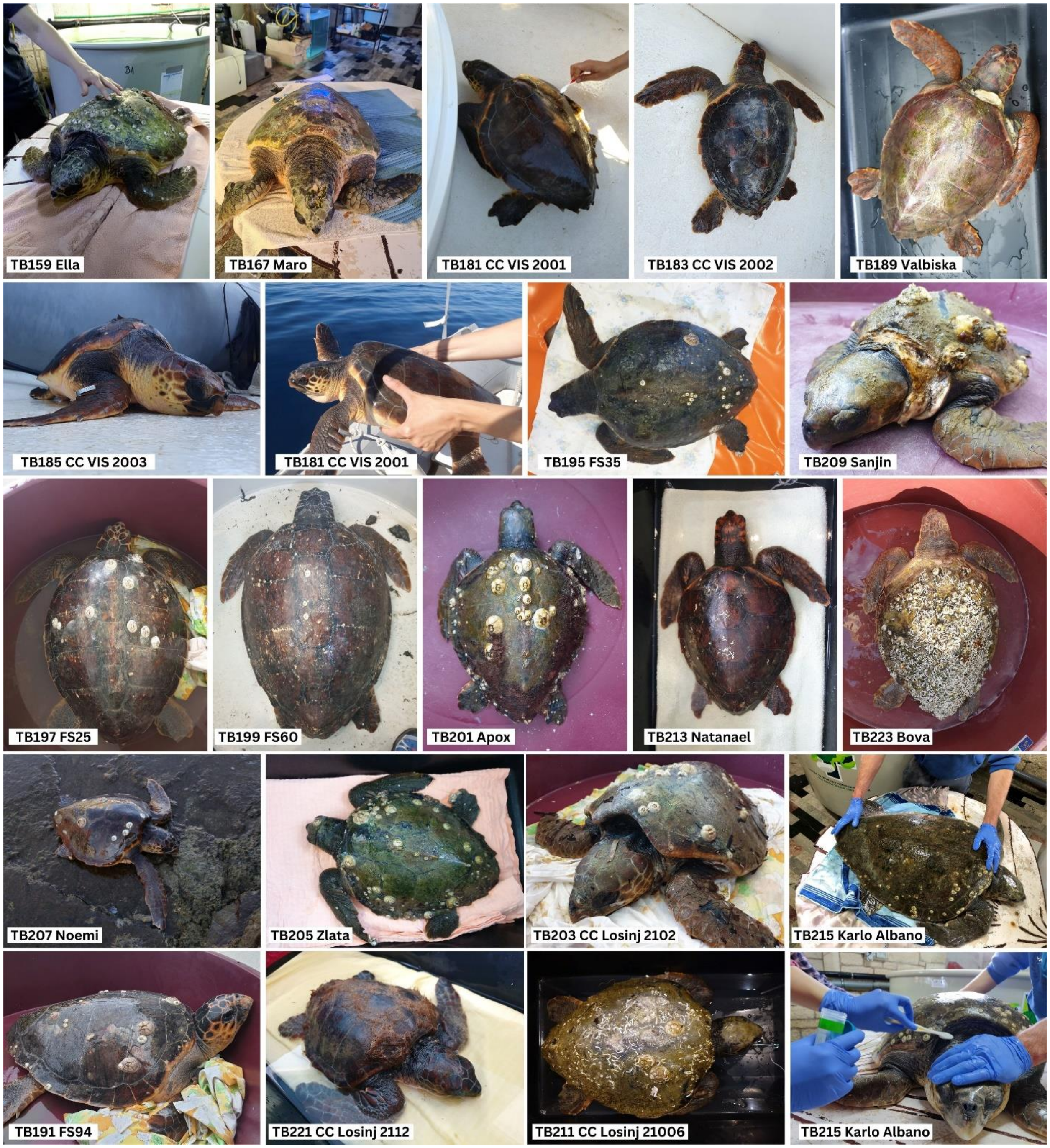


**Figure S1.** Photographs of sampled loggerhead sea turtles taken by Blue World Institute and Sea Turtle Rescue Centre Aquarium Pula; photographs of animals TB163, TB175, TB177, TB217, TB219, TB227, TB229 and TB231 were not taken.
