## Supplementary figures and images for "Growing older, growing more diverse: sea turtles and epibiotic cyanobacteria"

### Figure S2.pdf

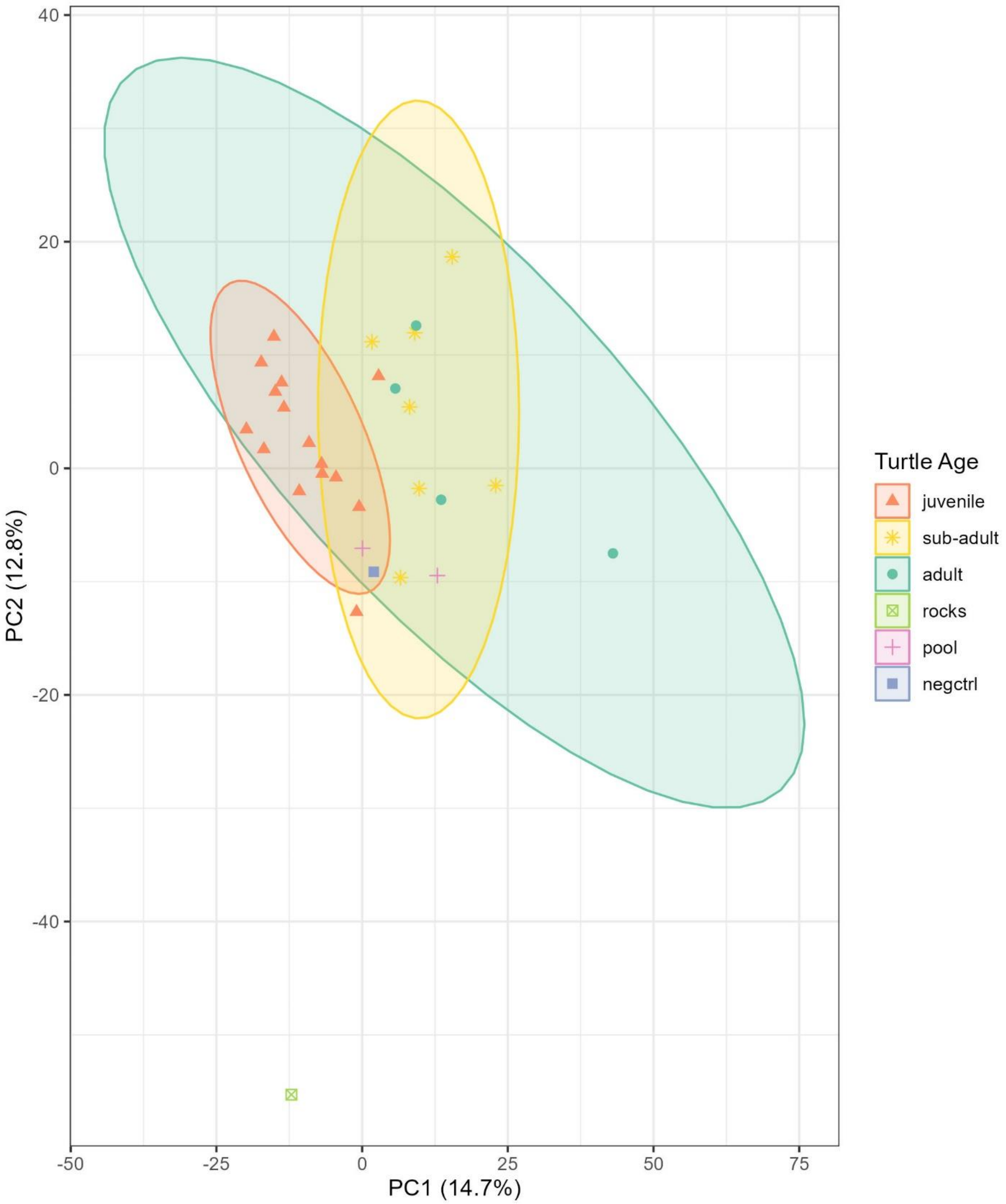


**Figure S2.** PCA of centar log ratio (CLR) transformed abundances for all samples.
